# Phosphorylation of AMPA receptor subunit GluA1 regulates clathrin-mediated receptor endocytosis

**DOI:** 10.1101/2020.12.03.410258

**Authors:** Matheus F. Sathler, Latika Khatri, Jessica P. Roberts, Regina C.C. Kubrusly, Edward B. Ziff, Seonil Kim

## Abstract

Synaptic strength is altered during synaptic plasticity by controlling the number of AMPA receptors (AMPARs) at excitatory synapses. In particular, during long-term potentiation and synaptic up-scaling, AMPARs are accumulated at synapses to increase synaptic strength. Neuronal activity leads to activity-dependent phosphorylation of AMPAR subunit GluA1, and subsequent increases in GluA1 surface expression, which can be achieved by either an increase in exocytosis or a decrease in endocytosis of the receptors. However, the molecular pathways underlying GluA1 phosphorylation-induced elevation of surface AMPAR expression are not completely understood. Here, we first employ fluorescence recovery after photobleaching (FRAP) to reveal that phosphorylation of GluA1 Serine 845 (S845) plays a more important role in receptor endocytosis than exocytosis during synaptic plasticity. Notably, endocytosis of AMPARs depends upon the clathrin adaptor, AP2, which recruits cargo proteins into endocytic clathrin coated pits. Importantly, the KRMK (Lysine-Arginine-Methionine-Lysine) motif in the carboxyl-terminus of GluA1 is suggested to be an AP2 binding site, but the exact function has not been defined. Moreover, the GluA1 KRMK motif is closely located to one of GluA1 phosphorylation sites, serine 845 (S845), and GluA1 S845 dephosphorylation is suggested to enhance endocytosis during long-term depression. In fact, we show that an increase in GluA1 S845 phosphorylation by two distinct forms of synaptic plasticity, long-term potentiation and synaptic up-scaling, diminishes the binding of the AP2 adaptor. This reduces endocytosis, resulting in elevation of GluA1 surface expression. We thus demonstrate a mechanism of GluA1 phosphorylation-regulated clathrin-mediated endocytosis of AMPARs.

## Introduction

Tetrameric α-Amino-3-hydroxyl-5-methyl-4-isoxazolepropionate-type glutamate receptors (AMPARs) mediate the majority of the fast excitatory synaptic transmission in the mammalian central nervous system (Diering and Huganir, 2018). AMPARs are formed by the assembly of subunits GluA1-4 in different combinations together with several auxiliary proteins that play critical roles in the receptor kinetics and trafficking patterns (Diering and Huganir, 2018; Greger et al., 2017; Henley and Wilkinson, 2016; Pick and Ziff, 2018). Regulation of AMPAR function is highly dynamic in many different forms of synaptic plasticity, including longterm potentiation and depression (LTP/LTD) and homeostatic synaptic plasticity (Diering and Huganir, 2018). A major part of these forms of synaptic plasticity is the trafficking of AMPARs from or to synapses to decrease or increase the number of AMPARs localized at synapses, which modulates the strength of synaptic activity (Hanley, 2018).

Although a large group of AMPAR auxiliary subunits can provide heterogeneity of AMPAR trafficking (Greger et al., 2017), activity-dependent receptor trafficking has long been known to be regulated by the phosphorylation of GluA1 mainly in a two-step process (Diering and Huganir, 2018; Pick and Ziff, 2018). First, phosphorylation of serine 845 (S845) in GluA1 is mediated by cAMP-dependent protein kinase A (PKA) or cGMP-dependent protein kinase II (cGKII) (Derkach et al., 2007; Serulle et al., 2007). Importantly, GluA1 S845 phosphorylation promotes GluA1 surface expression, increases channel open-probability, and mediates several forms of synaptic plasticity including LTP and synaptic up-scaling (Banke et al., 2000; Diering et al., 2014; Diering and Huganir, 2018; Ehlers, 2000; Esteban et al., 2003; Kim et al., 2015; Kim and Ziff, 2014; Lee et al., 2000; Lee et al., 2003; Man et al., 2007; Oh et al., 2006). In contrast, calcineurin-mediated dephosphorylation of S845 is involved in receptor endocytosis during LTD and synaptic down-scaling (Banke et al., 2000; Diering et al., 2014; Diering and Huganir, 2018;

Ehlers, 2000; Esteban et al., 2003; Kim et al., 2015; Kim and Ziff, 2014; Lee et al., 2000; Lee et al., 2003; Man et al., 2007; Oh et al., 2006). Second, when GluA1 is additionally phosphorylated at S831 by Ca^2+^/calmodulin-dependent protein kinase II (CaMKII) or protein kinase C (PKC), the single-channel conductance is elevated, contributing to the enhanced synaptic transmission following LTP induction, and GluA1-containing AMPARs are targeted to the postsynaptic density (PSD) (Banke et al., 2000; Barria et al., 1997; Derkach et al., 1999; Kristensen et al., 2011; Lee et al., 2000; Pick and Ziff, 2018). Therefore, cooperative phosphorylation on GluA1 plays important roles in AMPAR trafficking and function during synaptic plasticity.

Extensive studies have yielded inconsistent data on the role of GluA1 phosphorylation in AMPAR trafficking during synaptic plasticity. The impairment of synaptic plasticity in mice including by mutations that disrupt GluA1 phosphorylation strongly supports a requirement for receptor phosphorylation for synaptic plasticity (Lee et al., 2003). However, it has also been proposed that LTP expression does not require the carboxyl-tail of GluA1 (Granger et al., 2013). Furthermore, molecular mechanisms concerning the role of GluA1 phosphorylation in regulation of synaptic targeting and stabilization of AMPARs have not been fully understood. Importantly, LTP-inducing stimuli may promote AMPAR surface trafficking via S845 phosphorylation, supporting the idea that S845 phosphorylation has a direct role in GluA1-containing AMPAR exocytosis (Esteban et al., 2003; Lee et al., 2003). However, other studies reveal that LTD rather than LTP is strongly correlated with S845 dephosphorylation (Kameyama et al., 1998; Lee et al., 2000; Lee et al., 1998), indicating that S845 dephosphorylation is directly involved in LTD-induced endocytosis of GluA1-containing AMPARs (Lee et al., 2003). The latter idea is further supported by the finding that NMDA-induced AMPAR internalization correlates with the time of maximal dephosphorylation of S845, which is blocked by a calcineurin inhibitor (Ehlers, 2000). Moreover, it has been suggested that S845 phosphorylation stabilizes LTP by inhibiting internalization of recently inserted GluA1-containing receptors (Lee et al., 2003). Thus, GluA1 S845 phosphorylation may directly regulate AMPAR endocytosis rather than exocytosis, but the exact mechanisms underlying GluA1 S845 phosphorylation-mediated AMPAR endocytosis have not been investigated.

Activity-dependent endocytosis of AMPARs is mediated by the clathrin machinery (Ehlers, 2000; Hanley, 2018; Lee et al., 2002; Man et al., 2000; Parkinson and Hanley, 2018). Clathrin-mediated endocytosis (CME) is a multi-step process that requires several proteins to be recruited to specific membrane domains (Hanley, 2018; McMahon and Boucrot, 2011). The resulting protein complex changes the membrane geometry to create an invagination that leads to pit formation, ultimately resulting in scission of the formed vesicle from the plasma membrane by dynamin (McMahon and Boucrot, 2011). A central player in this process is an adapter protein complex, AP2, which binds to cargo proteins, endocytic accessory proteins, and clathrin (Fiuza et al., 2017; Kelly and Owen, 2011; Robinson, 2004; Traub, 2009). Significant progress has been made in identifying adaptor proteins responsible for regulating endocytosis of AMPARs (Hanley, 2018), however most of these proteins are GluA2-specific interactors (Fiuza et al., 2017; Hanley, 2018; Lee et al., 2002). For example, AP2 is known to bind to GluA2 at a KRMK (Lysine-Arginine-Methionine-Lysine) motif in the carboxyl terminus that is located proximal to the transmembrane domain, which promotes AMPAR endocytosis (Lee et al., 2002). Moreover, protein interacting with C-kinase 1 (PICK1), after interacting with the AP2 complex with GluA2, is recruited to clathrin-coated pits (Fiuza et al., 2017). This association is increased during NMDAR-mediated LTD and is able to promote dynamin activation, promoting GluA2 endocytosis (Fiuza et al., 2017). Interestingly, GluA1 is known to interact with the AP2 complex (Lee et al., 2002). More importantly, GluA1 contains the KRMK sequence, a candidate site for the AP2 complex binding, in the carboxyl terminus (Lee et al., 2002). The KRMK sequence in GluA1 is separated by 14 amino acids from S831 and by 29 amino acids from S845 (Diering and Huganir, 2018). Significantly, S-nitrosylation of cysteine 875 increases the binding of AP2 to GluA1 (Selvakumar et al., 2013), suggesting that changes in structure of distal sites on the carboxyl-terminus of GluA1 via post-translational modifications can affect AP2 complex binding and receptor internalization. Thus, it is possible that S845 phosphorylation may regulate the interaction between AP2 and GluA1. However, roles of the interaction between the KRMK sequence in GluA1 and the AP2 complex in AMPAR trafficking have not been completely investigated in depth. In particular, whether phosphorylation of GluA1 S845 can affect this interaction is not yet understood. Therefore, further clarification is needed to understand GluA1 endocytic mechanisms.

Here, we first employ fluorescence recovery after photobleaching (FRAP) to show that phosphorylation of GluA1 S845 has a more important role in endocytosis than in exocytosis. Moreover, we reveal that phosphorylation of GluA1 S845 promoted by two distinct pathways of synaptic plasticity, homeostatic up-scaling and chemically induced LTP (cLTP), is sufficient to decrease the endocytosis rate of GluA1 via the reduction of GluA1 binding to the AP2 complex. This ultimately leads to an increase in GluA1-containing AMPAR surface expression. Our results thus provide a molecular mechanism for how GluA1 S845 phosphorylation regulates GluA1-containing AMPAR trafficking.

## Results

### GluA1 S845 phosphorylation has no major role in receptor exocytosis in synapses

Increased AMPAR surface expression is mediated by either decreased endocytosis or increased exocytosis. To distinguish between these mechanisms, we examined by FRAP whether GluA1 S845 phosphorylation plays a critical role in surface insertion of AMPARs. To elucidate surface delivery of GluA1-containing AMPARs, we expressed super-ecliptic pHluorin (SEP)-tagged GluA1, which exhibits stronger fluorescence when exposed to pH 7.4 extracellular media and is nearly nonfluorescent when exposed to the acidic environment of endosomes (Yudowski et al., 2007). Using FRAP, we first determined what fraction of the SEP-GluA1 wild type (WT) signal on spines and dendritic shafts in DIV (days *in vitro)* 14 mouse cultured hippocampal neurons recovered under basal conditions or following chemical LTP (cLTP) (**Fig. 1**). Under basal conditions, we found approximately 65% SEP fluorescence recovery at 10 min after photobleaching, while significantly lower recovery was found after cLTP (GluA1 WT Basal, 66.40% ± 1.492 and GluA1 WT cLTP, 36.49 ± 1.103, p<0.0001) (**Fig. 1b-c**). The previous study has shown that SEP-GluA1 WT expressed in cultured neurons is largely mobile on spines in the absence of activity, which has more space available at the synapse to be occupied by newly trafficked GluA1 (Makino and Malinow, 2009). However, GluA1 stays much longer on spines and becomes immobile following LTP, thus there is much less space at the synapse for newly trafficked GluA1 (Makino and Malinow, 2009). Therefore, FRAP in the absence of LTP induction is significantly higher than the SEP recovery after LTP stimulation (Makino and Malinow, 2009). In fact, our data confirmed these findings that GluA1-containing AMPARs were incorporated into synapses and became immobile after cLTP induction. To elucidate the direct role of S845 phosphorylation on the exocytosis of GluA1, we generated a mutant GluA1, in which serine was replaced by alanine at the 845 position (S845A) of the carboxyl-terminus, which prevented GluA1 S845 from being phosphorylated (Lee, 2006; Lee et al., 2003). Under basal conditions, the SEP recovery of mutant GluA1 was slightly lower than GluA1 WT recovery after photobleaching (GluA1 WT Basal, 66.40% ± 1.492 and GluA1 S845A Basal, 60.39 ± 1.475, p=0.0203), while we found no significant difference in recovery kinetics (GluA1 WT Fast HalfLife = 3.265 ± 0.577 and GluA1 S845A Fast HalfLife = 3.568 ± 0.621, p=0.6147) (**Fig. 1b-c**). This suggests that phosphorylation of S845 may not be required for GluA1 constitutive exocytosis under basal conditions. We then compared SEP-GluA1 WT and SEP-GluA1 S845A recovery after cLTP induction. SEP-GluA1 S845A showed higher recovery rates when compared to SEP-GluA1 WT following cLTP induction (GluA1 WT cLTP, 36.49 ± 1.103 and GluA1 S845A cLTP, 42.34 ± 2.028, p=0.0157) (**Fig. 1b-c**). Moreover, mutant GluA1 reached a plateau much faster than GluA1 WT in the early phase of FRAP (GluA1 WT Fast HalfLife = 6.175 ± 1.733 and GluA1 S845A Fast HalfLife = 3.658 ± 1.262, p<0.0001). This indicates that a higher percentage of mutant receptors is mobile at synapses when compared to GluA1 WT-containing receptors, and thus GluA1 S845A-containing AMPARs are unable to be retained at the synapse following cLTP. These results thus suggest that phosphorylation of S845 plays important roles in GluA1 retention at the synaptic surface rather than receptor exocytosis following cLTP.

**FIGURE 1.**
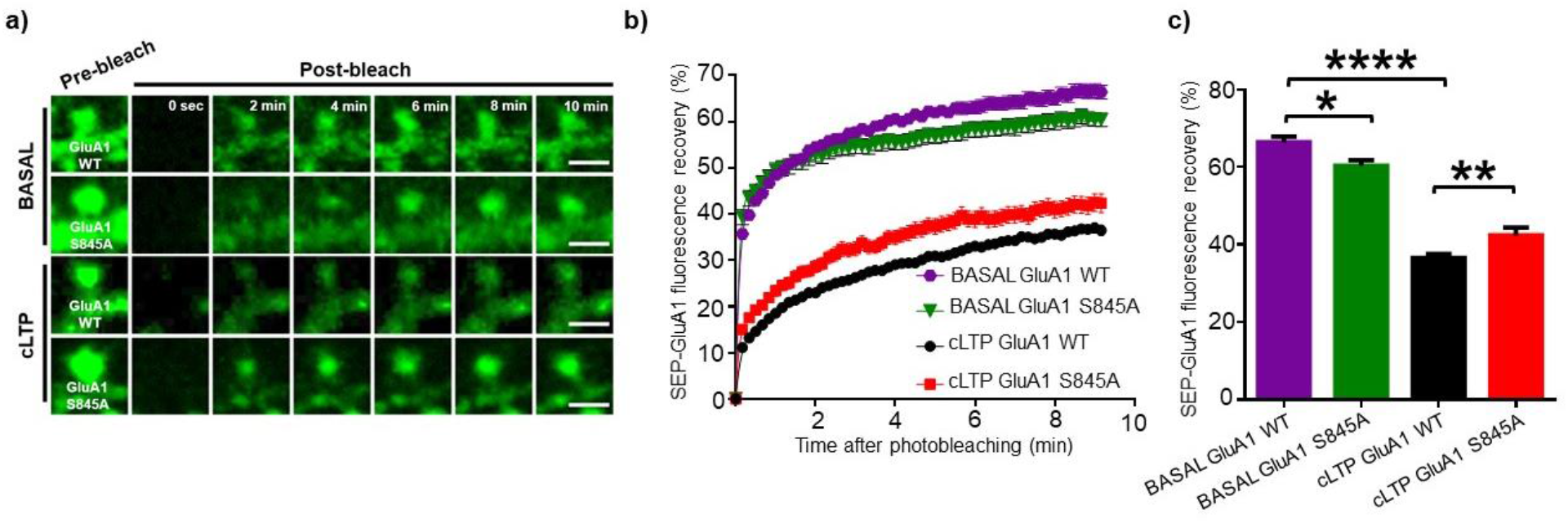
Phosphorylation of S845 is necessary for GluA1 synaptic retention rather than receptor exocytosis. **(a)** Representative images of fluorescence recovery after photobleaching (FRAP) experiments of SEP-GluA1 WT and SEP-GluA1 S845A under basal and cLTP conditions. Images depicted are before photobleaching, right after photobleaching, 2, 4, 6, 8 and 10 minutes after photobleaching for each condition. Images portrayed size are the exact photobleach ROI. Scale bar represents 5 μM. **(b)** Representative normalized traces of FRAP for each experimental group. Each point represents one image acquired every 10 seconds. Recovery rates were normalized to prephotobleaching intensities. Two phase association nonlinear fit was performed, and the fast halflife of each group was analyzed and compared with unpaired two-tailed Student’s t test. BASAL GluA1 WT and BASAL GluA1 S845A display similar recovery kinetics, while under cLTP, GluA1 S845A mutant display faster recovery than GluA1 WT, suggesting a higher percentage of mobile receptors. **(c)** Representative bar graphs of SEP fluorescence recovery at 10 minutes after photobleaching for each experimental group. BASAL GluA1 WT display a much higher recovery than cLTP GluA1, suggesting that under cLTP receptors are incorporated into synapses and become immobile. Under basal conditions, GluA1 S845 shows a small decrease in recovery when compared to GluA1 WT, although kinetics are similar. Under cLTP conditions, GluA1 S845A depicts a higher recovery rate than GluA1 WT, suggesting in the absence of S845 phosphorylation, receptors are unable to be retained at synapses. (BASAL GluA1 WT n=64 ROIs, BASAL GluA1 S845A n=60 ROIs, cLTP GluA1 WT n=139 ROIs, cLTP GluA1 S845A n=88 ROIs. Each ROI represents a spine and its adjacent dendrict shaft. * p<0.05, **p<0.01,****p<0.0001, One-Way Anova, uncorrected Fisher’s LSD)

### cLTP decreases AP2 adaptor binding to GluA1

Given that our FRAP data revealed that GluA1 S845A-containing receptors were more mobile at synapses following cLTP (**Fig. 1**), we explored the possibility that GluA1 S845 phosphorylation could regulate receptor endocytosis rather than exocytosis by examining interaction of GluA1 with the clathrin adaptor AP2, which has been reported to bind both GluA1 and GluA2 (Lee et al., 2002). First, co-immunoprecipitation (Co-IP) analysis with cell lysates of DIV14 cultured rat cortical neurons confirmed that β-Adaptin, the main subunit of the AP2 adaptor, was able to bind to endogenous GluA1, and this binding accumulated significantly when we blocked dynamin by treating cells with 1μM dynole to inhibit the scission of vesicles from the membrane (CTRL, 1.000 and Dynole,1.462 ± 0.1368, p=0.0279) (**Fig. 2a**). This confirmed that for GluA1 to be endocytosed. similar to GluA2, GluA1indeed interacted with β-Adaptin. To further examine whether activation of neurons to elevate GluA1 S845 phosphorylation and subsequently increase their surface expression was able to modify the binding of β-Adaptin to GluA1, we applied the same cLTP protocol used in the FRAP experiments (**Fig. 1**) and carried out co-IP experiments in DIV14 cultured neurons. We confirmed cLTP induced elevation of GluA1 S845 phosphorylation (CTRL, 1.000 and cLTP, 5.559 ± 0.8430, p=0.0002) (**Fig. 2b**) as seen previously (Diering et al., 2016). We further revealed a significant decrease in the binding of β-Adaptin to GluA1 in response to cLTP induction (CTRL 1.000 and cLTP 0.2403 ± 0.08304, p<0.0001) (**Fig. 2c**). These results suggest that LTP induction is able to increase the phosphorylation of S845, decreasing the interaction between GluA1 and the AP2 adaptor, which may contribute to LTP-induced elevation of GluA1 surface expression.

**FIGURE 2.**
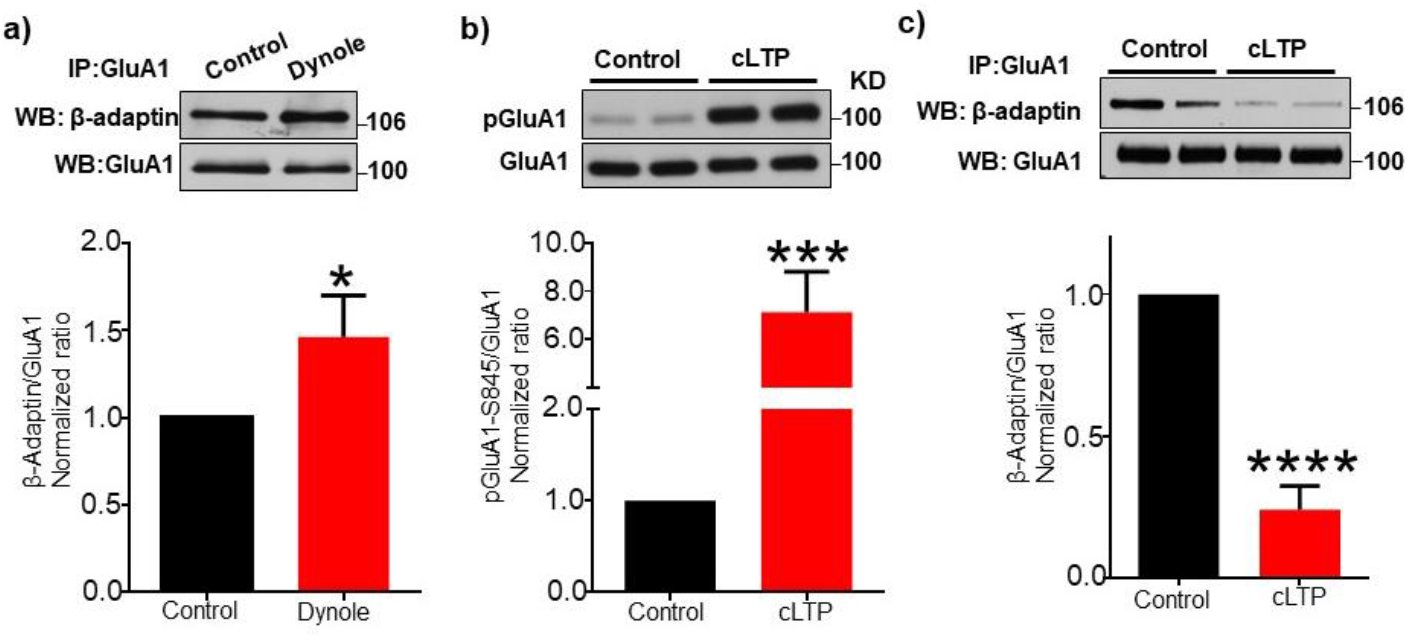
Chemical LTP increases phosphorylation of S845 and reduces binding of β-Adaptin to GluA1. **(a)** Representative immunoblots and quantitative analysis of co-immunoprecipitations of GluA1 from cultured cortical neurons showing that inhibition of dynamin by Dynole 1μM for 30 min increases the binding of β-Adaptin to GluA1. (n=3 different cultures, *p<0.05, unpaired two-tailed Student t tests). **(b)** Representative immunoblots and quantitative analysis of whole cell extracts from the same samples used in **(c)** for immunoprecipitation, showing that cLTP induction increases phosphorylation of S845. (n=4 different cultures, duplicated. ***p<0.001, unpaired two-tailed Student’s t test). **(c)** Representative immunoblots and quantitative analysis of co-Immunoprecipitation of GluA1 from cultured cortical neurons showing that cLTP induction decreases the amount of β-Adaptin bound to GluA1. (n=4 different cultures, duplicated. ****p<0.0001, unpaired two-tailed Student’s t test).

### TTX treatment-induced synaptic scaling up reduces the binding of GluA1 to the AP2 adaptor and reduces GluA1 endocytosis

We further explored the role of GluA1 phosphorylation and its interaction with the AP2 adaptor in another form of synaptic plasticity. While LTP increases the activity of individual synapses, during homeostatic synaptic plasticity there is a global increase in synaptic strength (Galanis and Vlachos, 2020). We induced synaptic up-scaling by chronically inhibiting neuronal activity, a well-established protocol to increase GluA1 S845 phosphorylation and subsequent surface expression (Diering et al., 2014; Kim and Ziff, 2014), and examined the effect on binding of GluA1 to β-Adaptin. We treated neurons with 2μM tetrodotoxin (TTX) for 48 hours and measured the interaction of GluA1 with β-Adaptin by using Co-IP. Our results showed that TTX treatment for 48 hours was sufficient to decrease the binding of β-Adaptin to GluA1 (CTRL, 1.000 and TTX, 0.4371 ± 0.1184, p=0.0034) (**Fig. 3a**). We also employed another protocol to elevate GluA1 S845 phosphorylation and surface expression by inhibiting calcineurin, a phosphatase that is known to dephosphorylate GluA1 and reduce surface GluA1 expression by promoting receptor endocytosis (D’Amelio et al., 2011; Kim and Ziff, 2014). We treated neurons with 5μM FK506 for 48 hours, a condition that increases GluA1 S845 phosphorylation and surface expression in cultured neurons (Kim and Ziff, 2014), and measured the interaction of GluA1 with β-Adaptin by using Co-IP. As expected, reduced calcineurin activity was capable of significantly decreasing β-Adaptin binding to GluA1 (CTRL, 1.000 and FK506, 0.4836 ± 0.08704, p=0.0051) (**Fig. 3a**). To examine whether reduced binding of β-Adaptin following synaptic up-scaling promoted reduced endocytosis of GluA1, we treated DIV 14 rat cultured hippocampal neurons with 2μM TTX for 48 hours and measured surface expression of GluA1 and GluA1 endocytosis rate (**Fig. 3b-c**). As shown previously (Diering et al., 2014; Kim and Ziff, 2014), TTX treatment was sufficient to increase GluA1 surface expression (CTRL, 1.000 and TTX, 1.175 ± 0.04437, p=0.0076) (**Fig. 3c**). By using an antibody live staining and feeding protocol to label the surface and endocytosed GluA1, we were able to show that GluA1 endocytosis rate was significantly decreased after TTX treatment (CTRL, 1.000 and TTX, 0.6306 ± 0.02549, p<0.0001) (**Fig. 3d**). These results confirmed that GluA1 surface accumulation following homeostatic up-scaling was mediated by reduced rates of GluA1 endocytosis. Taken together, these results suggest that during homeostatic up-scaling, an increase in S845 phosphorylation significantly reduces binding of the AP2 adaptor to GluA1, which decreases the endocytosis rate, ultimately contributing to increased surface accumulation of GluA1.

**FIGURE 3.**
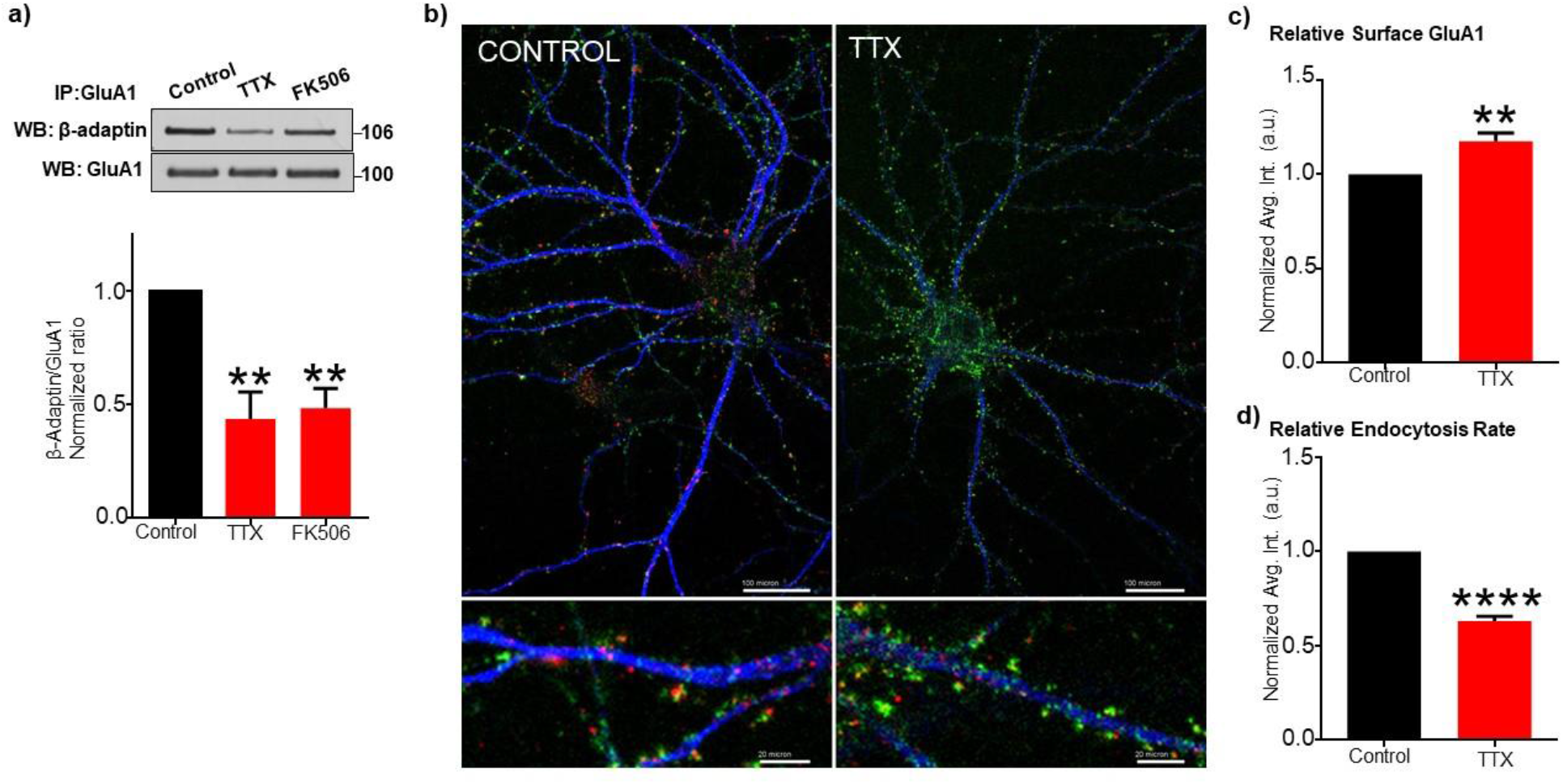
Scaling up protocols induce reduction of β-Adaptin binding to GluA1 and reduced endocytosis. **(a)** Representative immunoblots and quantitative analysis of co-Immunoprecipitations of GluA1 from cultured cortical neurons showing that TTX treatment (2μM for 48h) and FK506 treatment (5μM for 48h) decrease β-Adaptin binding to GluA1. (n=3 different cultures, **p<0.01, One-Way Anova, uncorrected Fisher’s LSD). **(b)** Neurons were treated with 2μM TTX for 48hr (+TTX) or left untreated (-TTX) and then incubated with a rabbit GluA1 extracellular domain antibody for 15 min at 37°C to allow the formation and endocytosis of antibody-GluA1 complexes. Antibody complexes remaining on the surface were removed by a brief acidic wash, neurons were fixed, and surface receptors relabeled with a mouse GluA1 extracellular domain antibody. Cells were permeabilized and surface and endocytosed antibody-GluA1 complexes were differentially stained with fluorescein- and rhodamine-labeled secondary antibodies (donkey) respectively. A scale bar indicated 100 μm (upper panel) and 20 μm (bottom panel). **c-d)** Quantification of receptor levels by confocal microscopy revealed that treatment of neurons for 48hr with TTX dramatically reduced GluA1 endocytosis and increased surface GluA1. n=4 different cultures, average of 10 neurons imaged per culture. **p<0.01 and **** p<0.0001, unpaired two-tailed Student t tests.

### Mutations on GluA1 CTD impact the binding of AP2

To examine the direct role of S845 phosphorylation on the binding of the AP2 adaptor to GluA1, we generated GST-fusion proteins containing the GluA1 cytoplasmic terminal domain (CTD) and various mutants. We incubated streptavidin beads conjugated with GST-fusion CTDs with brain cytosolic extracts in a GST pull-down assay. GST and GST-GluA2 CTD served respectively as negative and positive controls. Both the GluA2 and GluA1 CTD (GluA1C) were able to bind to β-Adaptin, while the GST control was unable to interact with β-Adaptin (GluA1C, 1.000, GST, 0.3046 ± 0.1195, p=0.0011, and GluA2, 1.690 ± 0.2652, p=0.0021) (**Fig. 4**), which was consistent with previous reports (Lee et al., 2002). We generated a S845D mutant in which serine was substituted for a phospho-mimetic aspartate (D) at position 845. Significantly, binding between GluA1 S845D and β-Adaptin was significant reduced (GluA1C, 1.000 and S845D, 0.3282 ± 0.07356, p=0.0015) (**Fig. 4**), confirming that GluA1 S845 phosphorylation was sufficient to reduce binding of the AP2 complex, resulting in a decrease in GluA1 endocytosis. Given that the KRMK motif in GluA2 is critical for AP2 interaction (Lee et al., 2002), we generated additional mutants of the KRMK motif in the GluA1 CTD (K819A, R820A, M821A and K822A) to determine if the KRMK motif in GluA1 was required for β-Adaptin binding. The GST-pulldown assay revealed that three mutants, including K819A, M821A and K822A, bound to β-Adaptin significantly less than GluA1C, whereas R820A was able to interact with β-Adaptin similarly to GluA1C (GluA1C, 1.000, K819A, 0.4785 ± 0.1946, p=0.0165, R820A, 0.7029 ± 0.2011, p=0.1167, M821A, 0.3253 ± 0.07782, p=0.0014, and K822A, 0.3981 ± 0.1080, p=0.0039) (**Fig. 4**), indicating that K819, M821 and K822 in GluA1 were required for AP2 binding, which was similar to GluA2.

**FIGURE 4.**
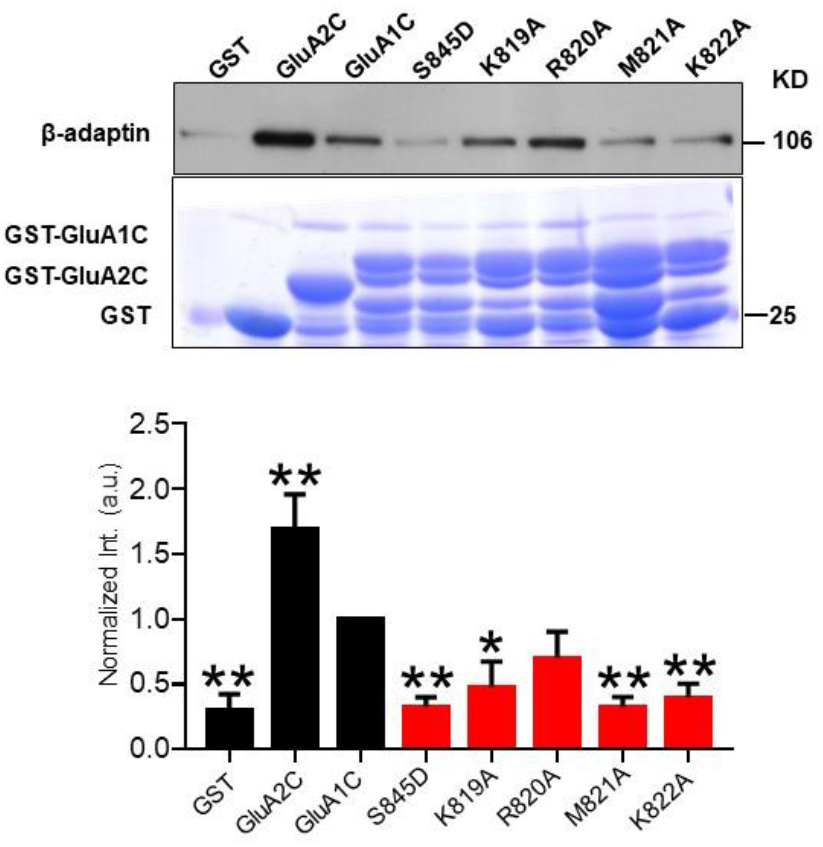
Interaction of β-Adaptin with GluA1 phosphomimetic and KRMK motif mutants is impaired. GST fusion proteins of the cytoplasmic tails of GluA1 (GluA1C) and its mutants were incubated with brain cytosolic extracts in a GST pull-down assay. Binding between phosphomimetic mutant GluA1 (S845D) and β-Adaptin was significantly reduced. Mutations in putative β-Adaptin binding sites (KRMK motif) also showed less binding to β-adaptin (n=4 different experiments. *p<0.05 **p<0.01, One-Way Anova, uncorrected Fisher’s LSD). GST and GluA2C were served as a negative and positive control for β-Adaptin pull-down, respectively. Commassie staining showed amount of GST-fusion proteins bound to beads in each lane.

### S845A mutant display more binding of AP2 to GluA1

We next examined the direct interaction of AP2 with GluA1 WT and GluA1 S845A in neurons. We infected DIV14-17 cultured cortical neurons with sindbis virus expressing HA-GluA1 WT or HA-GluA1 S845A for 16 hours and performed co-IP with an antibody against the HA tag. We observed that β-Adaptin interaction with HA-GluA1-S845A was significantly higher than HA-GluA1 WT (HA-GluA1 WT, 1.000 and HA-GluA1 S845A, 4.393 ± 1.480, p=0.0491) (**Fig. 5a**), further implicating S845 phosphorylation as a regulator of AP2 binding to GluA1. To further explore the role of S845 phosphorylation on the binding of AP2 to GluA1, we treated DIV14-17 cortical cultures with an activator or an inhibitor of PKA, a kinase that phosphorylate S845 (Diering and Huganir, 2018). We observed that blocking PKA activity with 1μM of KT5720 for 30 min was sufficient to increase the binding of β-Adaptin to GluA1 (**Fig. 5b**) (CTRL, 1.000 and KT5720, 1.484 ± 0.0052, p=0.0002), while PKA activation with 500μM 8-Br-cAMP decreased the binding of β-Adaptin to GluA1 (CTRL, 1.000 and 8-Br-cAMP 0.7217 ± 0.05915, p=0.0019). Thus, kinase activation during synaptic plasticity increases GluA1 S845 phosphorylation, which in turns decreases the interaction between GluA1 and β-Adaptin. This contributes to an elevation of AMPAR surface expression in response to synaptic plasticity.

**FIGURE 5.**
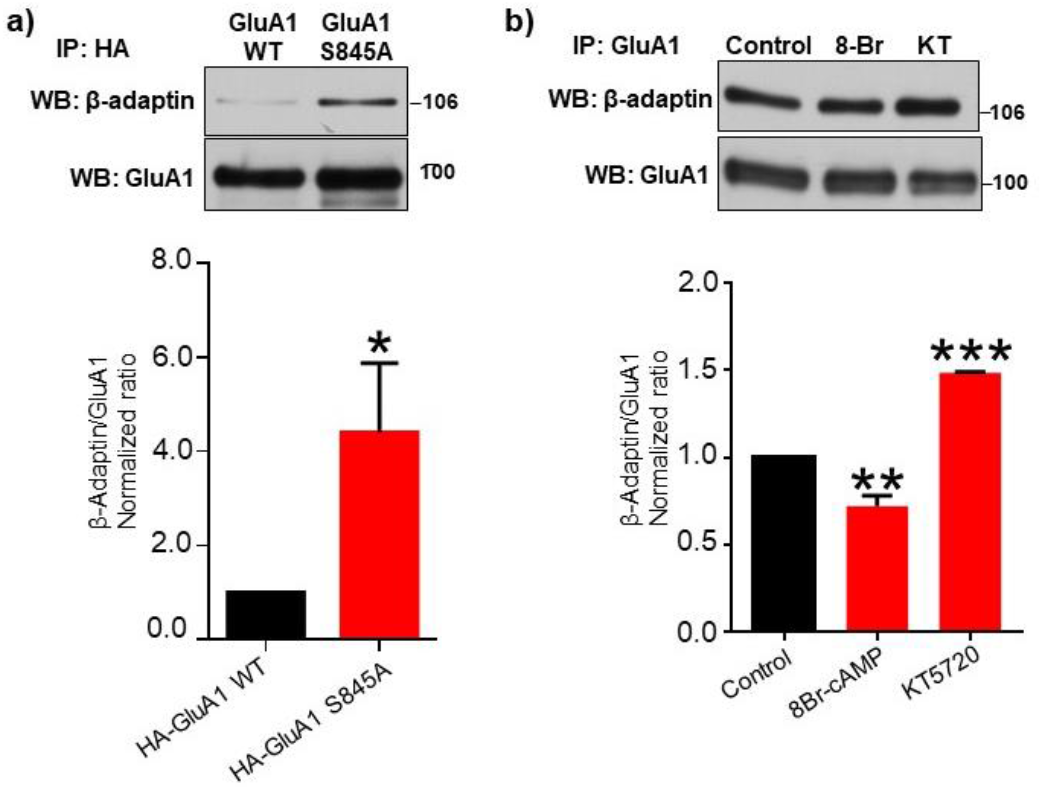
HA-GluA1-S845A receptors show less β-Adaptin binding to GluA1. **(a)** Representative immunoblots and quantitative analysis of co-Immunoprecipitation of HA-GluA1-WT and HA-GluA1-S845A virally expressed in cultured cortical neurons, showing that HA-GluA1-S845A mutant display more binding of β-Adaptin to GluA1. Overnight infection with HA-GluA1 WT and HA-GluA1-S845A was performed. (n=6 different cultures. *p<0.05, unpaired two-tailed Student t tests). **(b)** Representative immunoblots and quantitative analysis of co-Immunoprecipitation of GluA1 from cultured cortical neurons showing that treatment with PKA activator 8-Br-c-AMP (500μM for 10 min) decreases the binding of β-Adaptin to GluA1 while PKA blocker KT 5720 (1μM for 30 min) increases the binding of β-Adaptin to GluA1 (n=2-3 different cultures. **p<0.01, ***p<0.001. One-Way Anova, uncorrected Fisher’s LSD).

## Discussion

Synaptic plasticity is an activity-dependent alteration in synaptic strength, by means of control of the number of AMPARs at synapses (Park, 2018). In particular, hippocampal synaptic plasticity has long been considered a synaptic correlate for learning and memory (Park, 2018). GluA1-containing AMPARs have been suggested to play an important role in hippocampal synaptic plasticity including LTP and synaptic scaling (Diaz-Alonso et al., 2017; Diering et al., 2014; Hayashi et al., 2000; Jia et al., 1996; Kim and Ziff, 2014; Shi et al., 2001; Zamanillo et al., 1999). Importantly, the interplay between synaptic phosphorylation and dephosphorylation of GluA1 is central to regulation of AMPAR synaptic expression and synaptic plasticity (Diering and Huganir, 2018; Purkey and Dell’Acqua, 2020). During synaptic plasticity, synaptic strength can be enhanced by an increase in GluA1-containing AMPAR surface expression, which can be promoted by either increased exocytosis or decreased endocytosis of the receptors. However, the molecular mechanisms of how AMPAR trafficking is regulated by GluA1 phosphorylation have not been completely understood. Importantly, a previous study reveals that SEP-GluA1 is mobile on spines in the absence of activity (more turnover of AMPARs) (Makino and Malinow, 2009). However, following LTP, GluA1-continaing AMPARs are anchored at the synapse and becomes immobile at synapses following LTP, thus less newly trafficked GluA1 is incorporated at the synapse (Makino and Malinow, 2009). Consequently, FRAP of SEP-GluA1 WT is significantly lower after LTP induction, which is consistent with a model in which LTP brings and captures GluA1-containing AMPARs at synapses (Makino and Malinow, 2009). We also find less FRAP of SEP-GluA1 WT after LTP induction compared to FRAP without LTP stimulation, supporting this idea (**Fig. 1**). The mobility of GluA1-containing receptors from the dendrite to the extrasynaptic spine is likely to be rapid (Makino and Malinow, 2009). Thus, exocytosis of GluA1-containing AMPARs onto the dendrite during LTP also controls the GluA1 density at synapses (Makino and Malinow, 2009). Interestingly, FRAP of SEP-GluA1 S845A following cLTP induction is much higher and faster than SEP-GluA1 WT (**Fig. 1**), implicating that mutant GluA1 is more mobile than GluA1 WT and is less incorporated into synapses. Given that exocytosis and lateral diffusion of GluA1 S845A seem to be enhanced when compared to GluA1 WT, GluA1 S845 phosphorylation in fact plays a more important role in receptor endocytosis rather than exocytosis in response to synaptic plasticity, allowing receptors to be retained at synapses. Finally, under basal conditions, FRAP of mutant GluA1 at 10 min after photobleaching is slightly but significantly lower than FRAP of WT GluA1 (**Fig. 1**), which may also be due to less retention time of mutant GluA1 at synapses by accelerated endocytosis. Taken together, we determine that GluA1 S845 phosphorylation has no major role in receptor exocytosis, but in fact in preventing receptor endocytosis.

We used two distinct pathways of synaptic plasticity to enhance AMPAR synaptic strength by elevating GluA1 phosphorylation to reveal that GluA1 S845 phosphorylation prevents binding of the AP2 adaptor to receptors (**Fig. 2 and 3**), contributing to an increase in receptor surface retention. GluA1 endocytosis has been shown to be regulated by other forms of post-translational modifications of GluA1 CTD (Lin et al., 2009; Man et al., 2000; Selvakumar et al., 2013; Widagdo et al., 2015). Nitrosylation of GluA1 at C875 increases the binding of GluA1 to AP2 (Selvakumar et al., 2013). Moreover, palmitoylation of C811 prevents GluA1 interaction with protein 4.1N, a cytoskeletal scaffold protein, and reduces S818 phosphorylation by PKC (Diering and Huganir, 2018; Lin et al., 2009). In contrast, S818 PKC-mediated phosphorylation increases GluA1 interaction with 4.1N and limits receptor endocytosis (Diering and Huganir, 2018; Lin et al., 2009). Interestingly, ubiquitination of GluA1 is also able to regulate receptor endocytosis (Lin and Man, 2014; Widagdo et al., 2015). Ubiquitination of lysine 868 of GluA1 recruits epidermal growth factor receptor substrate 15 (Eps15) (Lin and Man, 2014), another endocytic adaptor of the clathrin pathway that is constitutively associated with AP2 (Benmerah et al., 1996; Benmerah et al., 1995; Lin and Man, 2014) and increases GluA1 internalization (Lin and Man, 2014). Interestingly, the GluA1 KRMK motif has two lysine residues available for ubiquitination, which raises the possibility that ubiquitination of these lysine residues (K819 and K822) can block AP2 interaction with GluA1. We also confirm that K819, M821 and K822 in GluA1 are required for AP2 binding (**Fig. 4**), which was similar to GluA2 (Lee et al., 2002).

The GluA1 CTD requirement for LTP was challenged by a report showing that LTP requires AMPAR trafficking, independent of subunit type (Granger et al., 2013). In addition, recent studies have demonstrated that the extracellular amino-terminal domains of AMPARs govern their trafficking for synaptic plasticity dependent on the AMPAR subunit types (Diaz-Alonso et al., 2017; Watson et al., 2017). Moreover, a study has reported that the levels of GluA1 phosphorylation are too low to regulate GluA1-dependent synaptic plasticity (Hosokawa et al., 2015). However, another study demonstrates that significantly higher levels of GluA1-containing AMPARs are phosphorylated at either S845 or S831 under basal conditions (Diering et al., 2016). Nonetheless, LTP expression has been thought to be dependent on the rapid synaptic insertion of GluA1-containing AMPARs (Hayashi et al., 2000; Shi et al., 2001), while endocytosis of AMPARs is important for LTD expression (Beattie et al., 2002; Carroll et al., 1999; Ehlers, 2000; Lee et al., 2003; Man et al., 2000). A study using phosphomutant (S831A and S845A) mice demonstrates that LTP is not completely absent, but LTD is completely dependent on GluA1 phosphorylation (Lee et al., 2003), supporting our idea that that GluA1 phosphorylation primarily regulates receptor endocytosis and is less important for exocytosis at excitatory synapses. Interestingly, PKA and calcineurin are targeted to GluA1 via the interaction with A-kinase anchoring protein 150 (AKAP150) in postsynaptic density of excitatory synapses, which regulate GluA1 phosphorylation (Sanderson et al., 2012). A study using mutant AKAP150 lacking calcineurin anchoring motif also shows an increase in GluA1 S845 phosphorylation and the inhibition of LTD (Sanderson et al., 2012), emphasizing the role of S845 phosphorylation in receptor endocytosis. Therefore, our results provide a molecular mechanism for how GluA1 S845 phosphorylation regulates GluA1-containing AMPAR clathrin-mediated endocytosis.

It has been shown that endocytic recycling endosomes containing AMPARs can serve as primary intracellular membrane compartments mobilized to the plasma membrane in response to LTP-inducing stimuli (Ehlers, 2000). Several proteins associated with recycling endosomes, including rab11, syntaxin 3, syntaxin 13, Eps15 homology domain protein Rme1/EHD1, synaptotagmin-1, synaptotagmin-7, complexin, SNAP-47, and synaptobrevin-2, have been reported to regulate AMPAR exocytosis during LTP (Ahmad et al., 2012; Grant et al., 2001; Jurado et al., 2013; Lin et al., 2001; Park et al., 2004; Prekeris et al., 1998; Reim et al., 2001; Ullrich et al., 1996; Wu et al., 2017; Zerial and McBride, 2001). However, it has not been shown that exocytosis dependent on these proteins is regulated by GluA1 phosphorylation, supporting our findings that the major impact of GluA1 phosphorylation is on receptor endocytosis during synaptic plasticity.

GluA1 homomeric AMPARs are Ca^2+^ permeable that plays important roles in several different types of synaptic plasticity, including homeostatic synaptic plasticity (Kim and Ziff, 2014), drug-related incubation of craving (Conrad et al., 2008) and NMDA receptor-mediated LTP (Barria et al., 1997; Diering and Huganir, 2018; Purkey and Dell’Acqua, 2020). However, the mechanism that enables the minority population of GluA2-lacking AMPA receptors to be accumulated at synapses during synaptic plasticity is not known. Interestingly, a study using mice specifically lacking phosphorylation of the GluA1 S845 site (GluA1 S845A mutant) demonstrates that this phosphorylation is required for GluA1 homomeric AMPAR surface retention (He et al., 2009). Specifically, in GluA1 S845A mutant neurons, homomeric receptors are removed from synapses mainly due to clathrin-mediated endocytosis and subsequent lysosomal degradation (He et al., 2009). Therefore, our findings further provide a mechanism of how GluA1 homomers are able to accumulate at synapses during synaptic plasticity.

GluA1 S845 phosphorylation is also implicated in several brain disorders (Zhang and Abdullah, 2013). Beta-amyloid peptide (Aβ) has been recognized as a causative factor for the cognitive impairments in Alzheimer’s disease (AD) and several studies have shown that Aβ leads to an increase in intracellular Ca^2+^ activity in cortical and hippocampal neurons (Brown et al., 2011; Busche et al., 2012; Busche et al., 2008; Harris et al., 2010; Hartley et al., 1999; Kuchibhotla et al., 2008; Liu et al., 2013; Minkeviciene et al., 2009; Palop et al., 2007; Palop and Mucke, 2009; Roberson et al., 2011; Sun et al., 2019; Verret et al., 2012). In particular, Aβ-induced Ca^2+^ hyperexcitation in hippocampal neurons stimulates Ca^2+^/calmodulin-dependent protein phosphatase, calcineurin, which dephosphorylates GluA1 S845 phosphorylation, enabling GluA1-containing AMPARs to be endocytosed from the plasma membrane (Lee et al., 1998; Sanderson et al., 2012). This leads to a reduction of synaptic strength and ultimately disrupts synaptic plasticity and cognitive function in AD (Forner et al., 2017). Another example is cocaine craving, a cue-induced cocaine seeking behavior that is intensified after withdrawal from chronic exposure of cocaine. The expression of incubated craving is mediated by GluA1 homomeric AMPARs in the nucleus accumbens (NAc) (Conrad et al., 2008). In fact, elevated synaptic levels of GluA1 homomeric receptors result in part from an increase in GluA1 S845 phosphorylation (Ferrario et al., 2011). Furthermore, enhanced GluA1 S845 phosphorylation and surface expression play important roles in the antidepressant effects of ketamine (Zhang et al., 2016). Taken together, GluA1 S845 phosphorylation has a unique role in various brain disorders and may serve a novel therapeutic target for these diseases.

## Methods

### 1) Expression Vectors

Plasmids and viral vectors expressing HA-GluA1, GST-R2C and GST-R1C have been described (Greger et al., 2003; Osten et al., 2000; Srivastava et al., 1998). GST-R1C mutants were cloned by PCR and ligated into pGEX-4T1 (Pharmacia). The HA-GluA1 S845A mutant was generated by the QuickChange mutagenesis kit (Stratagene). SEP-GluA1 WT plasmid was a gift from Dr Thomas Blanpied at University of Maryland School of Medicine and has been previously described (Kerr and Blanpied, 2012). SEP-GluA1-S845A was generated by PCR based QuikChange^®^ Site-Directed Mutagenesis Kit (Agilent) according to the manufacturer’s protocol. The following primers were used for mutagenesis 5-CTC-CCC-CGG-AAC-GCT-GGG-GCA-GGA-GCC-AGC-3 and 5-GCT-GGC-TCC-TGC-CCC-AGC-GTT-CCG-GGG-GAG-3. All mutants were confirmed by sequencing.

### 2) Antibodies

Purified Mouse Anti-Adaptin β Clone 74/Adaptin β (RUO) (BD Biosciences – 610381); Anti-Glutamate receptor 1 antibody (Millipore – AB1504); Anti-GluR1-NT (NT) Antibody, clone RH95 (Millipore – MAB2263); Anti-Glutamate Receptor 1 Antibody, phosphoSer 845 (Millipore – AB5849); Anti-GluR1 Antibody (Calbiochem-PC246, discontinued. Replaced by Millipore – ABN241); Purified anti-MAP2 Antibody (Biolegend – 822501, previously Covance catalog #PCK-554P).

### 3) Viral production and neuronal infection

BHK cells were electroporated with RNA of pSinRep5-HA-GluA1 or the C-terminal mutants and the helper DH (26S) according to Sindbis Expression System manual (Invitrogen) and as previously described (Osten et al., 2000). The pseudovirions-containing medium was collected after 24h, and the titer for each construct was tested empirically in neuronal cultures. For experimental expression, neurons were infected at 14–21 DIV with titer resulting in infection of 5%–10% of neurons (typically 5–20 ml of a-MEM virus stock diluted in 600 ml conditioned NB-B27 medium per 1 well of 6-well dish). The infectious medium was applied for 16 hr. Expression with no obvious adverse effects on morphology of the infected neurons was observed the next day. For immunoprecipitation (IP) experiments, neurons were cultured at 1 million cells per 6 cm dish and infected at 13 DIV with 60–100 ul of virus stock in 3 ml of conditioned NB-B27 medium for 16 hr.

### 4) Primary hippocampal and cortical neuronal culture

Primary rat hippocampal and cortical neuron cultures were prepared by a previously described protocol (Lu et al., 2014; Osten et al., 1998; Restituito et al., 2011). Animal experiments were conducted in compliance with the Institutional Animal Care and Use Committee at the New York University School of Medicine. The day before dissection, coverslips or 6cm Petri dishes were coated with poly-L-lysine in boric acid buffer at 37°C overnight. Before dissection, coverslips or dishes were washed twice with PBS and stored in the incubator ready for plating neurons. Primary hippocampal and cortical neuron cultures were obtained from E18-19 Sprague Dawley rat embryos. Pregnant rats were anesthetized with CO_2_, and embryos were then removed. Dissection was carried out in ice-cold PHG buffer (1× PBS, 10 mM HEPES, and 0.6% glucose, pH 7.35). After decapitation of the head, cortices and hippocampus were isolated under a dissection microscope in the sterile hood. Hippocampi and cortices were separately trypsinized in 1× trypsin for 15 min at 37 °C, washed 3 times in dissection buffer, and then resuspended in 5 ml of plating medium (minimal essential medium, 10% horse serum, 0.45% glucose, 1 mM pyruvate, 1% penicillin/streptomycin) warmed to 37°C. Hippocampi and cortices were triturated with a 5 ml of sterile pipette until the cell suspension appeared homogeneous, and cells were then counted with a hemocytometer. Cells were plated at a density of 120,000 per coverslip or 1,000,000 per 6-cm Petri dish in plating medium. 2–4h after plating, all media were removed and replaced with Neurobasal medium supplemented with B27 supplement (Invitrogen), glutamine (500 μM), and antibiotics. Every 4 days, half of the volume of medium remaining on the cells was removed and replaced with fresh Neurobasal medium. Anti-glia growth drug was usually added to cover slips into growth media after 8 DIV.

For FRAP experiments, mouse hippocampal neuron cultures were prepared as described previously (Sun et al., 2019; Sztukowski et al., 2018). Hippocampi were isolated from postnatal day 0 (P0) CD-1 mouse (Charles River) brain tissues and digested with 10U/mL papain (Worthington Biochemical Corp. Lakewood, NJ). Mouse hippocampal neurons were plated on poly lysine-coated glass bottom dishes (500,000 cells), transfected using Lipofectamine 2000 (Invitrogen) with 2 μg DNA of SEP-GluA1 WT or SEP-GluA1 S845A on DIV4, and imaged on day *in vitro* (DIV) 14. Cells were grown in Neurobasal Medium (Life Technologies, Carlsbad, CA) with B27 supplement (Life Technologies, Carlsbad, CA), 0.5mM Glutamax (Life Technologies) and 1% penicillin/streptomycin (Life Technologies). Colorado State University’s Institutional Animal Care and Use Committee reviewed and approved the animal care and protocol (16-6779A).

### 5) Neuronal Immunocytochemistry

Cells were initially treated with 2uM TTX for 48h before the experiment, as previously described (Pick et al., 2017). After drug treatment, 14 – 17 DIV hippocampi cultures were stained live with a GluA1 antibody (GluA1 Calbiochem PC246, rabbit, 1:20) for 15 min at 37°C in a humidified chamber, to allow for antibody binding and antibody/receptor complex internalization (endocytosed receptors). Cells were then washed with an acidic buffer (0.5M NaCl, 0.2M Acetic Acid, 1X PBS) for striping of remaining antibodies at cell surface, followed by three quick washes in PBS and fixed for 10 min at room temperature in a 4% PFA, 0.12M sucrose, PBS solution. Cells were then incubated with a different GluA1 antibody (Millipore – MAB2263, mouse, 1:500) to allow for staining of the remaining surface GluA1 (not endocytosed receptor). Cell were permeabilized for 5 min in 0.2% Triton X-100 and washed three times with PBS for 5 min at room temperature. Cells were blocked in 10% BSA in PBS for 1 h at room temperature and then incubated with a MAP2 antibody diluted in 3% BSA (MAP2 chicken, 1:10000, Biolegend) for 60 min at room temperature. Cells were washed three times in PBS with 5 min washes and incubated with secondary antibody diluted in 3% BSA for 60 min at room temperature bound to fluorophores (488 or 568 or 647) either from molecular probes (1:1,000) or from Jackson (1:300). Cells were then washed three times with PBS for 5 min, mounted on coverslips, and stored. Immunofluorescence images were acquired on a Nikon PCM 2000 confocal microscope under 60× objective. All images were acquired using the same settings for one experiment.

### 6) Fluorescence recovery after photobleaching (FRAP)

FRAP experiments were performed as previously described (Kim et al., 2016; Sole et al., 2019) using a spinning disk confocal microscope based on a Yokogawa CSUX1 system built on an Olympus IX83 inverted stand coupled to an Andor laser launch containing 405, 488, 561, and 637 nm diode lasers, 100–150 mW each. Images were collected using a 60× Plan Apo N 1.4 NA objective and two iXon EMCCD cameras (DU-897, Andor). This system is equipped with the ZDC constant focus system and a Tokai Hit chamber and objective heater. The stage and objective were heated to 37°C. Photobleaching was performed as previously described (Kim et al., 2016; Sole et al., 2019) using the FRAPPA system (Andor). Images were acquired every 10 s during 10 min, and the recovery of SEP-GluA1 WT and SEP-GluA1 S845A fluorescence within the bleached region (10 × 10 pixels) was quantitated as described below.

### 7) Image analysis and quantitation

Images were analyzed using ImageJ software. For immunocytochemistry, the areas of three dendrites on each image were manually defined by outlining the MAPII signal, and the intensity of the fluorescence of surface GluA1 or endocytosed GluA1 within the outlined dendrites to the total area was calculated for each dendrite and divided by total GluA1 intensity (surface GluA1 + endocytosed GluA1 fluorescence intensity). For FRAP, the average fluorescence intensity within each photobleached region was quantitated for each time point that images were acquired, background subtracted and normalized to the initial prephotobleach signal. All data was plotted into GraphPad Prism 9 and a two-phase association nonlinear fit was performed.

### 8) Co-Immunoprecipitation

Cells were initially treated with drugs, depending on the experiment and as described in each figure [TTX, Tocris Biosciences; FK506, Abcam; Dynole, Abcam; 8-Br-cAMP, Tocris Biosciences; KT 5720 (KT), Tocris Biosciences]. Co-immunoprecipitation was carried out as described previously (Sathler et al., 2016), with some modifications. Briefly, 14-17 DIV cortical cultures, were collected, homogenized in 1 ml of radioimmune precipitation assay buffer containing 50 mM Tris, pH 7.4, 150 mM NaCl, 1% Nonidet P-40, 0.5% deoxycholate, 0.1% SDS, 10 mM EGTA, 10 mM EDTA, phosphatase inhibitor cocktails I and II (Sigma), and Protease Inhibitor Complete (Roche Applied Science, Indianapolis, IN, USA). For input sample, 50 uL of protein lysate was removed, rocked for 60 min at 4 C, and boiled in an equal volume of 2X loading buffer. The immunoprecipitation was performed by incubating the protein lysate with 1ug of GluA1 antibody (Millipore, ab1504) or 1ug HA antibody (Santa Cruz) for 16 h at 4 C followed by binding to Protein A Plus-agarose beads (Santa Cruz) at 4C for 60 min. Beads were then pelleted by centrifugation and washed 3 times in wash buffer (50 mM Tris, pH 7.4, 300 mM NaCl, 5 mM EGTA, 0.1% Triton), suspended in 1X loading buffer (30 ul), and boiled (immunoprecipitated sample). Equal volumes were loaded for Western blotting, and all membranes were probed with anti-beta-adaptin antibody (1:1000, BD Transduction) and GluA1 or HA antibody to control for amount precipitated.

### 9) GST Pulldown

GST-fusion proteins were expressed and purified as described (Osten et al., 1998). GST, GST-R2C,GST-R1C or GST-R1C mutants (10 ug each) bound on Glutathione Sepharose (Pharmacia) beads were incubated with whole brain cytosolic fraction for 3 hours at 4C. Beads were then pelleted by centrifugation and washed 3 times in wash buffer (20 mM HEPES, 150 mM NaCl, 0.1% Triton), suspended in 1X loading buffer (30 uL), boiled and equal volumes were loaded for western blotting. The bottom half of the acrylamide gel was stained with Commassie Blue and the GST stained bands were used as loading control.

### 10) Chemical LTP protocol

cLTP protocol was followed as previously described (Zheng et al., 2015). Briefly, DIV 14–17 cortical cultured neurons were washed in Mg^2+^ free buffer (NaCl 150 mM, CaCl_2_ 2 mM, KCl 5 mM, HEPES 10 mM, glucose 30 mM, strychnine 1 μM, bicuculline 20 μM) 3 times, and incubated in glycine buffer (Mg^2+^ free buffer with 0.2 mM glycine) at 37°C for 15 min. Then, Mg^2+^ buffer (Mg^2+^ free buffer with 2 mM MgCl_2_) was added to block NMDARs and cells were incubated at 37°C for 30 min before being processed for co-immunoprecipitation or FRAP.

### 11) Statistics

All statistical comparisons were analyzed with the GraphPad Prism9 software. Unpaired two-tailed Student’s t tests were used in single comparisons. For multiple comparisons, we used one-way ANOVA followed by Fisher’s Least Significant Difference (LSD) test to determine statistical significance. Results were represented as a mean ± SEM, and a p value < 0.05 was considered statistically significant.

## Acknowledgements

We thank Joseph Pick, Danielle Ferreira, and the rest of the Ziff Lab and Kim Lab for their helpful discussion and comments. We also thank Dr Michael Tamkun and the Tamkun Lab for their help with the FRAP experiments. This work was supported by the Brazilian agency Conselho Nacional de Desenvolvimento Científico e Tecnológico (CNPq) (RCCK), NIH grant (R37AG013620 to EBZ), the pilot program from the Colorado Clinical and Translational Sciences Institute (SK), College Research Council Shared Research Program from Colorado State University (SK), and the Boettcher Foundation (SK)

